# High capacity and dynamic accessibility in associative memory networks with context-dependent neuronal and synaptic gating

**DOI:** 10.1101/2020.01.08.898528

**Authors:** William F. Podlaski, Everton J. Agnes, Tim P. Vogels

**Affiliations:** Centre for Neural Circuits and Behaviour, University of Oxford, OX1 3SR Oxford, United Kingdom and Champalimaud Neuroscience Programme, Champalimaud Foundation, 1400-038 Lisbon, Portugal; Centre for Neural Circuits and Behaviour, University of Oxford, OX1 3SR Oxford, United Kingdom and Biozentrum, University of Basel, 4056 Basel, Switzerland; Centre for Neural Circuits and Behaviour, University of Oxford, OX1 3SR Oxford, United Kingdom and Institute of Science and Technology Austria, 3400 Klosterneuburg, Austria

## Abstract

Biological memory is known to be flexible — memory formation and recall depend on factors such as the behavioral context of the organism. However, this property is often ignored in associative memory models. Here, we bring this dynamic nature of memory to the fore by introducing a novel model of associative memory, which we refer to as the context-modular memory network. In our model, stored memory patterns are associated to one of several background network states, or contexts. Memories are accessible when their corresponding context is active, and are otherwise inaccessible. Context modulates the effective network connectivity by imposing a specific configuration of neuronal and synaptic gating – gated neurons (respectively synapses) have their activity (respectively weights) momentarily silenced, thereby reducing interference from memories belonging to other contexts. Memory patterns are randomly and independently chosen, while neuronal and synaptic gates may be selected randomly or optimized through a process of contextual synaptic refinement. Through signal-to-noise and mean field analyses, we show that context-modular memory networks can exhibit substantially increased memory capacity with random neuronal gating, but not with random synaptic gating. For contextual synaptic refinement, we devise a method in which synapses are gated off for a given context if they destabilize the memory patterns in that context, drastically improving memory capacity. Notably, synaptic refinement allows for patterns to be accessible in multiple contexts, stabilizing memory patterns even for weight matrices that do not contain any information about the memory patterns such as Gaussian random matrices. Lastly, we show that context modulates the relative stability of accessible versus inaccessible memories, thereby confirming that contextual control acts as a mechanism to temporarily hide or reveal particular memories. Overall, our model integrates recent ideas about context-dependent memory organization with classic associative memory models, highlights an intriguing trade-off between memory capacity and accessibility, and carries important implications for the understanding of biological memory storage and recall in the brain.

---

Discrete attractor networks offer a simple and powerful model for auto-associative memory, enabling the retrieval of stored, content-addressable memory patterns from noisy external cues^1–3^. The biological and computational relevance of such associative memory models depends on the statistics of the memory patterns and the choice of connectivity architecture^2,4,5^, among other factors. While classic models typically feature uncorrelated patterns (i.e., half of the network units are active for each pattern, chosen uniformly at random) and all-to-all connectivity^1^, many other variants have been proposed and explored^2^. For example, sparse^6^, hierarchical^7,8^, and structured^9^ patterns have been shown to confer computational advantages to storage capacity and recall, and are more biologically plausible^10^. Furthermore, networks with sparse^11^ or modular^12–15^ connectivity exhibit improved storage ability and their structure may resemble brain circuits more closely^16,17^.

A common feature of much theoretical work on associative memory is the focus on memory capacity – the maximum number of stable patterns that the network is able to store^2,6,18,19^. While capacity is certainly a crucial quantity when considering the function of associative memory models, recent work in psychology and neuroscience has highlighted the flexibility of memory as an equally significant property. Memory *accessibility* – the relative ease by which a particular pattern can be retrieved – is not constant, and has been experimentally shown to vary over time depending upon the background context^20–25^. Such work has put forth the idea that the ability to flexibly store^26^ and recall^24^ memories depending upon the behavioral context may be a crucial aspect of memory’s functional role. Classic models of associative memory typically feature static and uniform memory stability, and thus not only fail in capturing this crucial aspect of biological memory, but also miss out on possible computational advantages of such an approach.

Here, we introduce a new model of associative memory, which we refer to as the *context-modular memory network.* In the model, retrieval dynamics are controlled by an externally-imposed background state, which we refer to as *context*. This control is exerted onto the network by silencing neurons and synapses, termed *neuronal gating* and *synaptic gating,* respectively. These gating schemes, which are defined in Section 1, modify the effective network connectivity, and thereby also the accessibility of memories, i.e., their stability. Based on signal-to-noise analysis and mean-field theory, we show in Section 2 that random neuronal gating improves memory capacity considerably when compared to the classic Hopfield model, while random synaptic gating gating has only a minimal increase in memory capacity. We then introduce the concept of *refinement* in Section 3, in which synaptic gating (but not neuronal gating) can be optimized to improve memory stability. We show that memory capacity is drastically increased with synaptic refinement. Finally, we demonstrate the model’s ability to stabilize memories in the active context and destabilize memories in the inactive context in Section 4, highlighting the model’s powerful control over memory accessibility. Overall, this work incorporates novel ideas about the dynamic and behaviorally-relevant use of memory into classic associative memory models, and may serve as a novel framework for the study of flexible associative memory.

## 1. Model

### 1.1. Model description

We introduce a context-dependent auto-associative memory network: the context-modular memory network. The main functionality of this model is to have dynamic memory accessibility — that is, depending on a background context signal, the model stabilizes a subset of memory patterns while keeping others hidden. This can be achieved by temporarily modifying the network such that there is a different effective connectivity matrix for each contextual state (Fig. 1(a)). To implement such a model, we modify a standard, fully-connected Hopfield network with *N* neurons^1^ to include *s* contextual states, i.e., contexts, which control two types of gating. First, in neuronal (nrn) gating, each context has a corresponding set of allocated neurons, *N*_cxt_, while the other *N* − *N*_cxt_ neurons are gated off. We use the parameter *a* = *N*_cxt_/*N* to denote the proportion of allocated neurons per context. Second, in synaptic (syn) gating, each contextual state defines a particular set of allocated connections (synapses) per neuron, *K*, with the other *N*−*K* connections being gated off. We use the parameter *c* = *K*/*N*_cxt_ to denote the connection density.

**Fig. 1:**
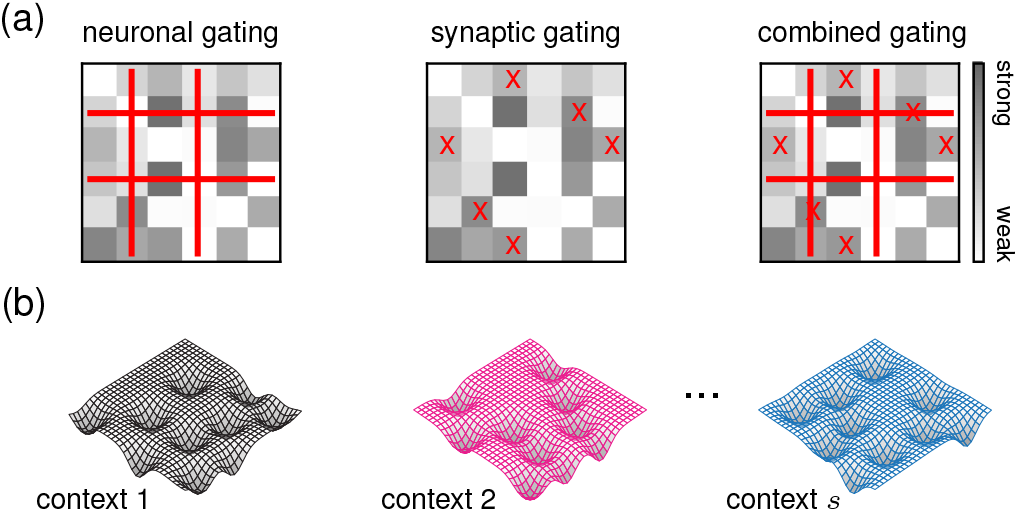
The context-modular memory network. (a) Contextual configurations alter the effective connectivity matrix of the associative memory network: neuronal gating removes particular columns and rows (left), synaptic gating removes individual elements (center), and together, they implement both effects (right). (b) Each contextual state produces a distinct energy landscape with a unique set of attractor memory patterns, with others remaining hidden.

We impose that only one context is active at any given time, and we consider the recall dynamics of the network from the perspective of this active context, which we denote by *k*. Units are denoted 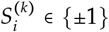 for neuron *i* when context *k* is active, with dynamics

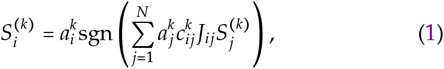

where *J_ij_* is the synaptic weight between neurons *j* and *i* (independent of context), and 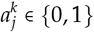 and 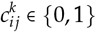 represent the effects of neuronal and synaptic gating for context *k*, respectively. We note that standard Hopfield dynamics correspond to setting 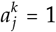 and 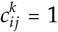 for all *i* and *j*. The network stores *p* memory patterns per context, making *P* = *sp* total memories (except for the case in which memories are randomly allocated to multiple contexts, explored in Section 3.3). Memory patterns are defined as 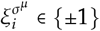 for neuron *i* and memory *μ* of context 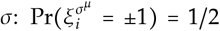. In the following sections, we explore connectivity matrices based on variants of Hebbian learning rules^1,6^.

We note that neuronal gating does not set neurons to the typical inactive state of the Hopfield network 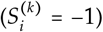, but instead to zero. This can be interpreted as setting all the output connections of the gated neuron to zero, but is equivalent to inactivation of the neuron in a more intuitive {0,1} formulation of the network (see Appendix A).

### 1.2. Model interpretation

The gating schemes defined above can be interpreted as mechanisms that impose a distinct effective connectivity matrix for each contextual state. The network dynamics can then be viewed as following a context-dependent energy function (Fig. 1(b)),

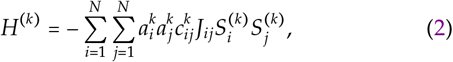

with context *k* being active. Neuronal gating temporarily removes particular rows and columns from the connectivity matrix (Fig. 1(a), left), whereas synaptic gating temporarily removes individual entries (Fig. 1(a), middle). The combination of the two produces a smaller and sparser connectivity matrix (Fig. 1(a), right), with a potentially large number of synaptic connections remaining hidden, to be used in other contexts. This enables the fully-connected network to exhibit an effectively modular structure, in which the active context *k* imposes an *inhibitory mask,* given by 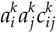, onto the connectivity matrix. It is thus intuitive to analyze the model in terms of the proportion of gated neurons, i.e., neuronal gating ratio, 1 − *a*, and proportion of gated synapses, i.e., synaptic gating ratio, 1 − *c*. The neuronal gating ratio indicates the proportion of neurons that are gated off while a given context is active. The synaptic gating ratio, usually referred to as dilution^27^, indicates the proportion of gated off synapses during recall of memories in the active context.

### 1.3. Memory capacity definitions

We define parameters related to pattern storage to study the maximum number of memories that can be successfully retrieved. The network *load* refers to the ratio of stored patterns to network size. Here, we have two notions of load. First, the *single-context load* is defined as

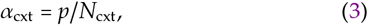

which is the number of patterns stored in each individual context divided by the number of allocated neurons per context. For simplicity, we impose that all contexts store the same number of patterns and have the same number of allocated neurons, and so the single-context load is identical across all contexts. Second, the *total load* is

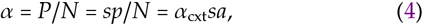

which is the total number of patterns stored in the network across all contexts, divided by the network size. The maximum load for which all patterns are stable attractors is known as the memory capacity. We refer to the single-context and total capacities as 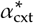 and *α*\*, respectively^28^. Due to analytical tractability, memory capacity is probed conditioned on an externally-imposed context without considering how context is implemented (see Discussion).

## 2. Random neuronal and synaptic gating

First we consider the case in which neurons and synapses are allocated uniformly at random to each context. That is, for each context *k*, variables 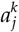 and 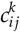 (Eq. 1) are set to 1 or 0 with probabilities 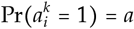 and 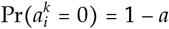 for neuronal gating and 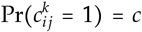 and 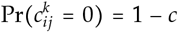 for synaptic gating. For simplicity and mathematical tractability, the synaptic gating matrix is symmetric; 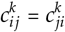. These quantities are used to define the weight matrix through a modified Hebbian rule as

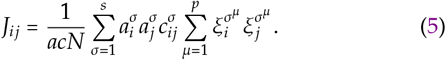

Thus, synaptic weights only contain information about memory patterns for contexts in which that synapse, and the corresponding pair of neurons, are allocated. Below, we calculate the capacity of these networks conditioned on an active background context with respect to the parameters *a*, *c*, and *s*. We use two established heuristic methods: signal-to-noise analysis and mean-field theory.

### 2.1. Signal-to-noise analysis

We consider the general case of a network with *s* contexts, and with neuronal and synaptic gating (both random), determined by the parameters *a* and *c* defined above. We calculate memory capacity following a signal-to-noise analysis^2^ that estimates the probability that a given neuron is in the incorrect state when recalling a memory pattern. To calculate this probability, we start by defining 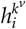 as the total input to neuron *i* when the network configuration is set to pattern *v* of context *k*. From this definition, we then calculate the probability that 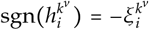 considering that neuron *i* is allocated to context *k*, i.e., 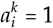. Combining Eqs. 1 and 5, we obtain

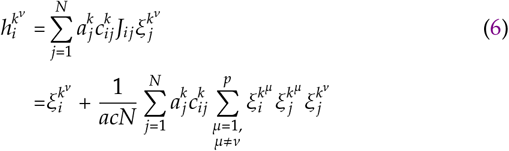

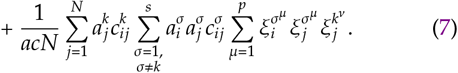

The first term on the right-hand side of Eq. 7 is the memory pattern whose stability is being evaluated. The second and third terms on the right-hand side of Eq. 7 represent two sources of crosstalk noise that may disrupt the stability of stored memories. The second term is analogous to the noise term in a standard Hopfield network^2^, accounting for the noise introduced by other patterns stored in the same context. The third term accounts for the noise introduced by patterns stored in different contexts. We multiply Eq. 7 by 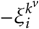, obtaining

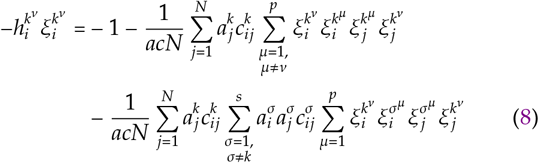

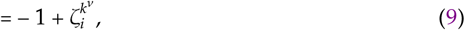

where 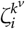 represents the combined effect of the two crosstalk terms multiplied by 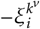,

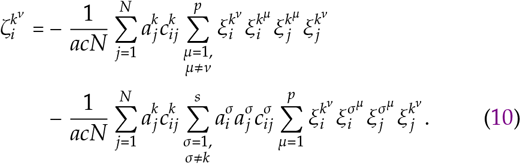

From Eq. 9 we note that if 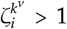, the input to neuron *i* has the opposite of the desired sign. To estimate memory capacity, we can thus calculate the probability of neuron *i* being in the incorrect state for memory *v* in context *k*, 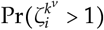.

We observe that, due the effects of the gating variables, both terms on the right-hand side of Eq. 10 are sums of discrete random variables, {−1,0,1}. Considering a sequence of Bernoulli random variables *X_j_* such that Pr(*X_j_* = 0) = Pr(*X_j_* = 1) = 1/2, we can approximate 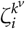 as

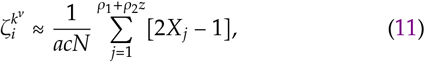

with the limits of the sum defined as

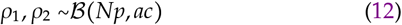

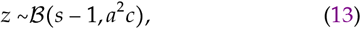

where 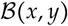 is a binomial distribution with *x* trials and probability of success *y*. Here, we can intuitively interpret *ρ*_1_ and *ρ*_2_ as the number of non-zero terms in the sums over *j* and *μ*, and *z* as the number of non-zero terms in the sum over *σ* in Eq. 10. In the thermodynamic limit, we can assume that *ρ*_1_ and *ρ*_2_ (Eq. 12) are well approximated by Gaussian noise. However, since the number of contexts *s* is finite, *z* is not well approximated by a Gaussian and must be treated differently. We thus assume that for pattern *v* of contextk, the realization of *z* takes a value 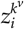 for neuron *i*. With this assumption, 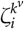 can be well-approximated as a Gaussian random variable, and using Wald’s equation (Appendix B) we obtain

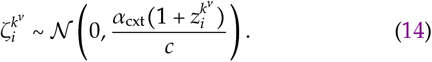

Here we take a shortcut by noting a correspondence between kV

the noise term 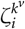 and previous results for the standard and diluted Hopfield networks^2,18,27^. Capacity should be reached when the variance approaches that of the standard Hopfield (for neuronal gating alone, *c* = 1) or diluted Hopfield (for synaptic gating, *c* < 1) networks. In those classic models, the crosstalk variance is approximately 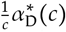 (Appendix C), where 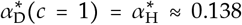. We thus define an equality between the variance of Eq. 14 and the diluted Hopfield variance. By rearranging terms and averaging over all neurons, patterns and contexts, we obtain a single-context capacity of

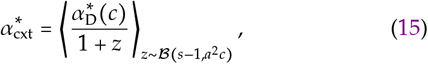

where 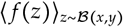 denotes an average over the binomial random variable *z*. The denominator of Eq. 15 can be interpreted as the effective number of contexts whose patterns contribute to the crosstalk noise — the number 1 refers to the currently-active context, which always contributes, and *z* refers to the contribution from other contexts, which depends on the parameters *s*, *a*, and *c*. In other words, each context can be viewed as a standalone diluted Hopfield network of size *N*_cxt_ storing *p* patterns, but with additional noise in the weights due to the influence of effectively *zp* other memories. The denominator 1 + *z* is always greater than or equal to 1, and so the single-context capacity is upper bounded by the diluted Hopfield network limit 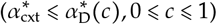.

From Eq. 15, we obtain a total capacity

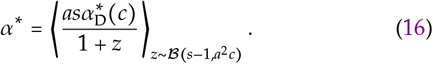

Eqs. 15 and 16 apply generally for cases of arbitrary numbers of contexts, *s*, proportion of gated neurons, 1 − *a*, and proportion of gated synapses, 1 − *c*. We make several observations about the signal-to-noise estimates of single-context capacity, 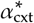, and total capacity, *α*\* (Fig. 2). Single-context capacity is upper-bounded by the Hopfield limit (Fig. 2(a,b)). For fixed proportion of gated neurons or synapses, the single-context capacity decreases as the number of contexts, *s*, increases (Fig. 2(a,b)). We observe that for high proportions of gated neurons (1 − *a* ≤ 0.75 and *c* = 1), single-context capacity degrades gradually (Fig. 2(a)), leading to large total capacity (Fig. 2(c)). For synaptic gating alone (*a* = 1), single-context capacity is very small (Fig. 2(b)) which leads to only a modest improvement in total capacity (Fig. 2(d)).

**Fig. 2:**
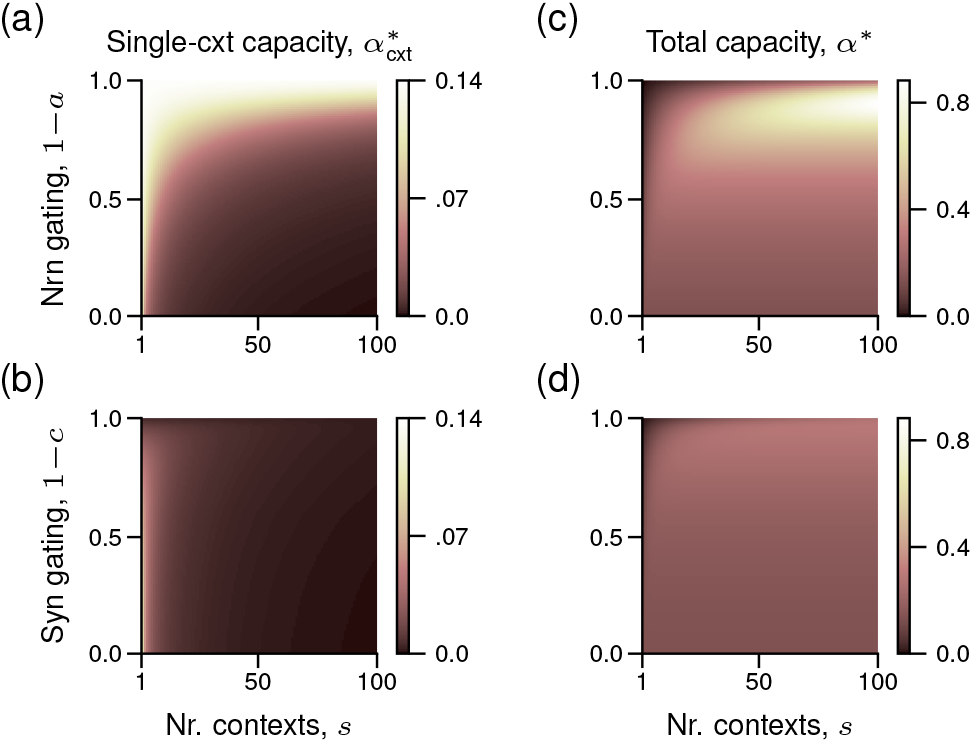
Memory capacity of the context-modular memory network with random neuronal and synaptic gating. (a) Capacity estimation for single-context capacity, 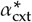, as a function of the number of contexts, *s*, and neuronal gating, 1 − *a*, for neuronal gating alone (*c* = 1) from Eq. 15. (b) Capacity estimation for single-context capacity,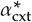, as a function of the number of contexts, *s*, and synaptic gating, 1 − *c*, for synaptic gating along (*a* = 1) from Eq. 15. (c,d) Total capacity for the networks considered in panels (a,b) based on Eq. 16.

Memory capacity is well described by Eq. 16, but it cannot provide a direct intuition of how different parameters affect capacity. To arrive at such an intuition, we consider the simplified case in which each neuron is gated in the same number of contexts. This is equivalent to replacing the variable *z* with its expected value in Eqs. 15 and 16. Doing so, we arrive at a single-context capacity given by

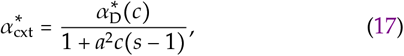

and a total capacity of

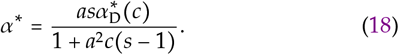

We can make several observations from these simplified equations. First, the optimal neuronal gating ratio, 1 − *a*_opt_, for a fixed number of contexts and no dilution (*c* = 1) is 1 − *a*_opt_ = 1 − s^−1/2^ (Fig. 3(a), red line). Furthermore, taking the limit of large *s* (such that 1 + *a*^2^(*s* − 1) ≈ *a*^2^*s*), the total capacity approaches *α*\* ≈ *α*_H_/*a* (Fig. 3(b), red line), and thus 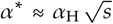 considering optimal neuronal gating. It follows that the high capacity emerges due to the patterns having low activity, growing sub-linearly as a function of the number of contexts. The optimal synaptic gating ratio, 1 − *c*_opt_, cannot be easily calculated in our simplified analysis because *α*_D_(*c*) does not have a closed-form solution^27^. However, 1 − *c*_opt_ ∝ 1 − *s*^−3/4^ fits well the optimal synaptic gating ratio (Fig. 3(a), purple line). Taking the limit of large *s* for synaptic gating alone (*a* =1), the total capacity approaches *α*\* ≈ *α*_D_(*c*)/*c* (Fig. 3(b), purple line). We also compared the results to a low-activity variant of the Hopfield network^6^ (Fig. 3(b), dashed gray line; Appendix). While the low-activity capacity is larger, it is only achieved for very sparse activity levels compared to neuronal gating. Overall, we see that the context-modular memory network uses contextual gating to harnesses sparsity in activity and connectivity in a similar way to previous models.

**Fig. 3:**
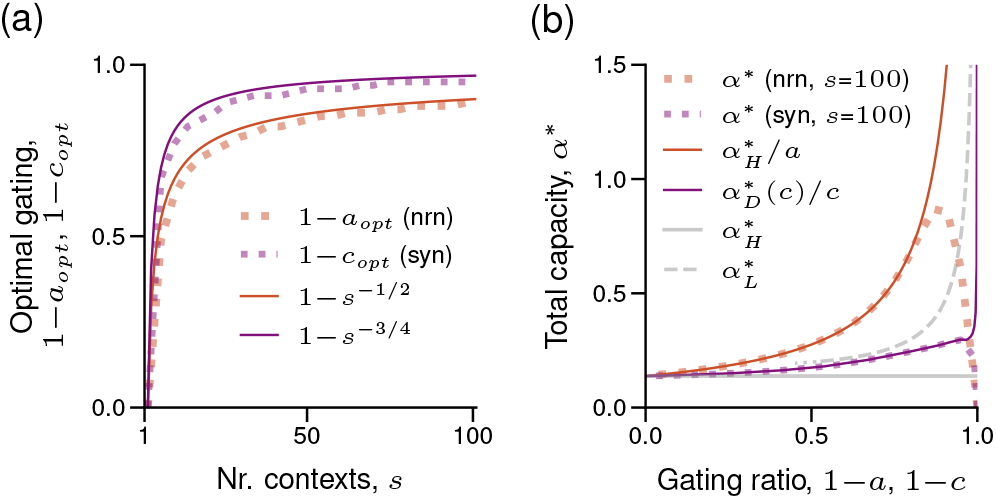
Optimal gating parameter and simplified total capacity. (a) Optimal neuronal gating ratio, 1 − *a*_opt_, for neuronal gating alone (*c* = 1, red), and optimal synaptic gating ratio, 1 − *c*_opt_, for synaptic gating alone (*a* = 1, purple), as a function of the number of contexts, *s*. Dashed lines are the optimal values, numerically estimated from Eqs. 15 and 16. Full lines are the functions indicated in the legend. (b) Total capacity, *α*\*, as a function of neuronal and synaptic gating ratios, 1 − *a* and 1 − *c*, respectively. Dashed thick lines are the estimated capacity for a fixed number of contexts (s =100) as a function of the neuronal gating ratio, 1 − *a*, for synaptic gating alone (*c* =1, red), and synaptic gating ratio, 1 − *c*, for synaptic gating alone (*a* = 1, purple) from Eq. 15. Red and purple full lines are the simplified estimated capacity with functions indicated in the legend. Full gray line is the Hopfield limit, 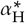, and dashed gray line is the estimated capacity of the low-activity Hopfield network, 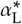 (Appendix C).

### 2.2. Mean-field theory

To validate our signal-to-noise results, we confirmed them using a heuristic mean-field approach^2^. We consider a stochastic version of the Hopfield network, with dynamics

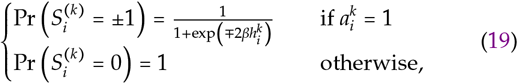

where 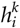 is the input to neuron *i* and *β* = 1/*T* is the inverse of the temperature, *T*. We replace the fluctuating input 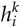 in Eq. 19 by its average value 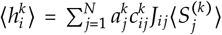 (where 〈·〉) represents thermal average), to obtain the average activation

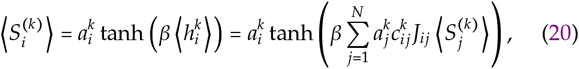

if context *k* is active.

Based on the mean-field theory (see Appendix D for detailed calculation), we arrive at the following three equations for the context-modular memory network,

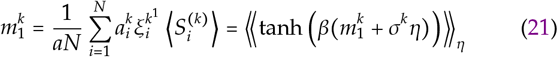

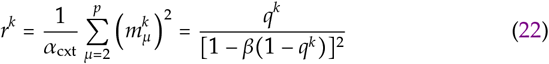

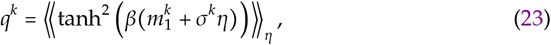

where 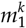 is the *overlap* between the average network state 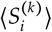 and pattern 1 of the active context *k*; *r^k^* is the *mean square overlap* of the system configuration with the non-retrieved patterns in context *k*; *q^k^* is the Edwards-Anderson order parameter^29,30^; 《·》_*η*_ represents a Gaussian average over the variable 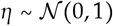; and *σ^k^* represents the crosstalk noise on pattern 1 of context *k* due to all other patterns. Then, writing 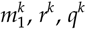, *r^k^*, *q^k^* and *σ^k^* as *m*, *r*, *q* and *σ*, respectively, we estimate the noise as

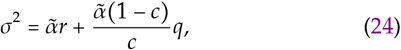

where 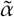 is given by

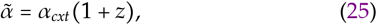

and 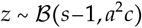 is a binomial random variable (analogous to Eq. 13). For expository purposes, we can consider the expected value for *z* to obtain

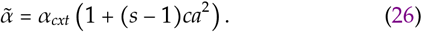

We observe that setting *a*, *c*, and *s* to 1 recovers the original Hopfield network capacity with *σ*^2^ = *α*_H_*r*. Furthermore, for *s* = 1 and *a* = 1, we can recover the diluted Hopfield capacity, with *σ*^2^ = *α*_D_*r* + *α*_D_(1 − *c*)*q*/*c*. We solve the mean-field equations in the zero temperature limit (*β* → ∞)^2^, obtaining precisely the same memory capacity as Eq. 16, thereby validating the signal-to-noise analysis. While the signal-to-noise analysis usually requires an arbitrary noise cutoff value^2^, we overcame this by directly comparing to the mean-field results for the standard and diluted Hopfield networks, 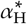 and 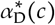.

Importantly, the mean-field theory can provide an estimate of the overlap between the stored memory patterns and the attractor states of the dynamics at capacity — such a measure is needed for numerical estimations of capacity. The level of overlap is well captured by the standard Hopfield network when there is no dilution (*c* = 1), and it’s well described by the levels of dilution otherwise^27^ (Appendix C). We can thus write the mean-field overlap *m*(*c*, *a*, *s*) as a function of the diluted network result, as

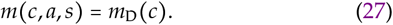

For neuronal gating alone (*c* = 1), the overlap between memory patterns and the fixed-points of the network’s dynamics is independent of *a* and *s* (*m*\* ≈ 0.97; Fig. 4, red line and dots)^30^. For synaptic gating alone (*a* = 1), the overlap is independent of *s*, but decreases with increasing dilution, 1 - *c* (Fig. 4, purple line and dots)^27^. This indicates that there is increased error in recall for large amounts of dilution (*c* ≪ 1).

**Fig. 4:**
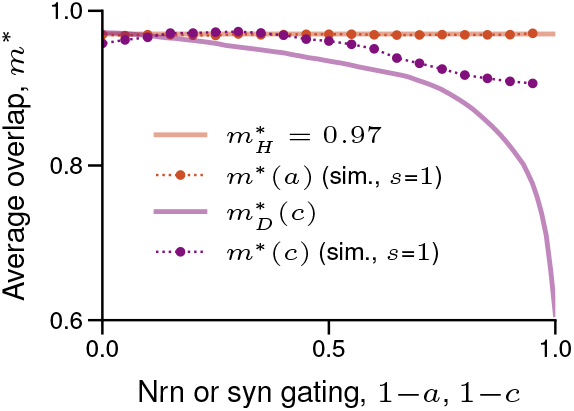
Mean-field overlap. Average mean-field overlap between memory patterns and fixed-points of the network’s dynamics for neuronal gating alone (*c* = 1, red) and synaptic gating alone (*a* = 1, purple) as a function of neuronal and synaptic gating ratios, 1 − *a* and 1 − *c*, respectively. Full lines indicate theoretical results from mean-field theory. Circles connected by dotted lines indicate results from simulations for networks at theoretical capacity for a single context (*s* = 1). See Appendix G for details.

To validate our analyses, we compared the theoretical results with numerical simulations of size *N* = 10,000 (Figs. 4, 5; simulation details can be found in Appendix G). We first verified the accuracy of the theoretical overlap, *m*\*, by simulating networks at theoretical capacity and measuring the average overlap. Simulations of neuronal gating alone showed overlap values very close to the theoretical prediction (Fig. 4, red points). However, while the simulations of synaptic gating alone resulted in a decreased overlap as the synaptic gating ratio, 1 − *c*, increased, it was substantially higher than expected, and highlights the difficulties in numerically simulating such diluted networks.

As for memory capacity, simulations and theory behave similarly for neuronal gating alone (*c* = 1; Fig. 5(a,b)), with discrepancies likely due to finite-size effects, and the approximations in the analysis described above. For synaptic gating alone (*a* = 1), there is an increased discrepancy between simulation and theory (Fig. 5(c,d)), again due to finite-size effects, but also due to the difficulty in achieving an accurate overlap cutoff, as mentioned above (Fig. 4).

**Fig. 5:**
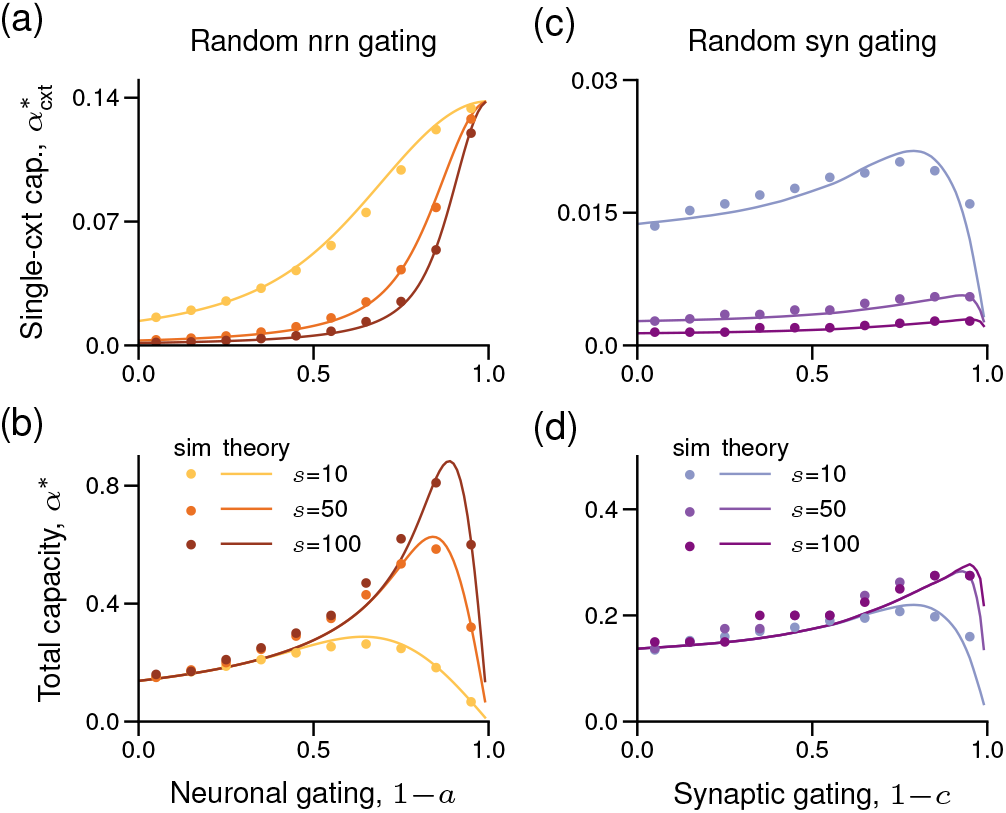
Comparison between mean-field theory and simulations for random neuronal and synaptic gating. (a,b) Mean-field capacity estimation (solid lines) and numerical simulations (dots) for single-context capacity, 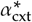 (panel (a)) from Eq. 15, and total capacity, *α*\* (panel (b)) from Eq. 16, as a function of neuronal gating ratio, 1 − *a*, for neuronal gating alone (*c* = 1). Plots for three different number of context, *s* = 10, *s* = 50, and *s* = 100. (c,d) Same as panels (a,b) but as a function of synaptic gating ratio, 1 − *c*, for synaptic gating alone (*a* = 1). *N* = 10,000 for all simulations (Appendix G).

### 2.3. Information content

While high storage capacity is a relevant and important aspect of an associative memory network, the information stored per pattern is also an essential aspect, as both should be maximized. The information content can be calculated as the total entropy (Shannon information) across all retrieved patterns^19^. This calculation is straightforward for neuronal gating, because the average overlap between the minima of the dynamics (retrieved patterns) and the original patterns is close to unity, *m* ≈ 0.97 (Fig. 4, red line). Due to the low values of overlap for synaptic gating (Fig. 4), we calculate the information content as the total entropy considering the original memory patterns minus the *missing information* due to inaccurate recall^31^. Based on the above definition, we obtain the following information content of the context-modular network (see Appendix E for detailed calculation),

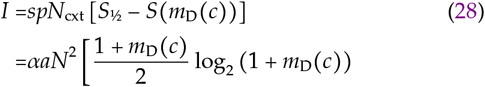

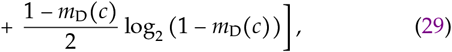

where *S*_½_ is the entropy per neuron given that 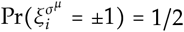, and *S*(*m*) is the missing information given that the average overlap between original memory patterns and corresponding attractors is *m*^31^. In the following, we consider information content relative to that of the standard Hopfield network (Appendix E).

The information content of the context-modular memory network with random neuronal gating alone (*c* = 1) and *s* = 100 contexts, when at maximum capacity (*α* = *α*\* in Eq. 29), follows closely that of the standard Hopfield network at maximum capacity (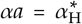 and 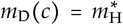 in Eq. 29) for small gating ratios 1 − *a* < 0.75 (Fig. 6, red). For larger gating ratios, we observe that the information content drops off steeply to zero. Similarly to storage capacity (Fig. 3(b)), this discrepancy is due to the dependency on the number of contexts, *s*, and suggests that a similar information content may be obtained for an arbitrarily high gating ratio provided that *s* is large enough. The information content of a network with random synaptic gating alone (*a* = 1) and *s* = 100 contexts is similar in magnitude to that of the standard Hopfield network, and grows larger with increased gating ratio, 1 − *c* (Fig. 6, purple). Analogously to neuronal gating, the information content for synaptic gating drops to zero as the gating ratio approaches 1, due to a finite number of contexts *s*.

**Fig. 6:**
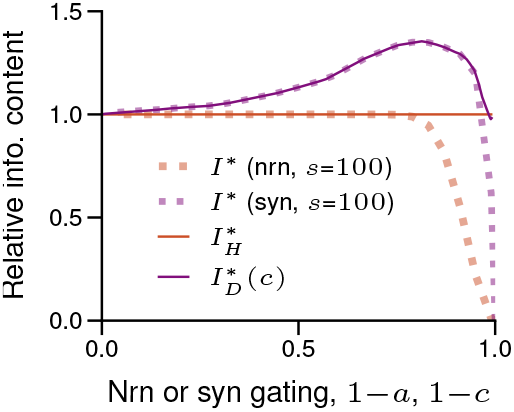
Information content. Relative information content of the context-modular memory network with random neuronal and synaptic gating. Dashed lines shows the information content of random neuronal gating alone (*c* = 1 and *s* = 100, red) and random synaptic gating alone (*a* = 1 and *s* = 100, purple) as a function of neuronal gating parameter, 1 − *a*, and synaptic gating parameter, 1 − *c*, normalized by the information content of the standard Hopfield network (solid red line). Solid purple shows the information content of the diluted Hopfield network as a function of connection density, 1 − *c*, normalized by the information content of the standard Hopfield network.

Overall, we see that while storage capacity grows large for context-modular memory networks with high gating ratios 1 − *a* and 1 − *c* (Fig. 3(b), red and purple lines), information content remains similar to the standard Hopfield network (Fig. 6). This is similar to the classic low-activity variant^6^, whose storage capacity diverges as activity levels approach zero (Fig. 3(b) dashed grey line), but whose information content remains finite (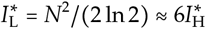^6,19^).

## 3. Synaptic refinement

We have shown that random synaptic gating does not provide considerable improvements to memory capacity. We thus devise a more selective way of imposing synaptic control which we refer to as synaptic *refinement*. Connections are refined after learning of memory patterns, so we consider that synaptic gating can only act on the recall dynamics (Eq. 1) and not on the weight matrix, *J_ij_*. We thus define the weight matrix based on the standard Hebbian learning rule (including random neuronal gating),

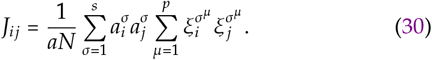

To intuitively describe the refinement scheme, we first consider the case of no neuronal gating (*a* = 1). In this case, Eq. 30 sets the synaptic weight between each pair of neurons according to the correlation in their activity across all patterns, acting to stabilize the majority of patterns^2^. However, the set of memories in the active context may produce substantially different correlations, such that some synaptic weights decrease the stability of the majority of memories in this active set. Such synaptic weights have a net harmful effect on memory recall, and performance would improve if these weights were gated off. This reasoning leads to a well-defined criterion for synaptic gating: if there is a mismatch between the sign of the synaptic weight serving to stabilize all memories versus serving to stabilize memories belonging to a specific context, then this synapse is gated off for this context. We then observe that this criterion can be combined with the inclusion of random neuronal gating (*a* < 1) before learning without one affecting the other.

More precisely, we find an optimal set of synaptic gates 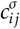 for stability of memory patterns in each context. First, we define a hypothetical “isolated” weight matrix for each individual context, e.g., context σ, with elements

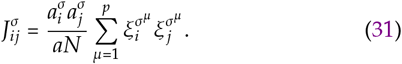

Notice that Eq. 31 only incorporates patterns from context σ, such that 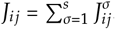. Then, the optimized gating state of each synapse is defined as

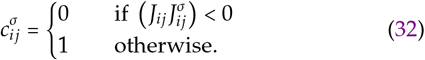

This method ensures that for context *σ*, the sign of each ungated weight reflects the sign of the correlation between neurons *i* and *j* over patterns in context *σ*, but not over all other patterns. Due to the symmetry in the weight matrices *J_ij_* and 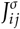, synaptic gating is symmetric 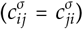. Furthermore, and importantly, the synaptic gating ratio, 1 − *c*, is no longer a predetermined parameter of the network, but is a function of the number of contexts, *s*, and the neuronal gating ratio, 1 − *a*.

### 3.1. Synaptic gating ratio

To estimate the expected number of gated synapses as a function of network parameters *s* and *a*, we assume that the overall weight distributions from Eqs. 30 and 31 are Gaussian with zero mean and variances proportional to the amount of crosstalk that they contribute, which is determined by a binomial random variable 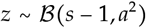 analogous to the signal-to-noise analysis from Section 2.1. This leads to synaptic gating given by (see Appendix F for detailed calculation)

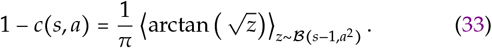

For synaptic refinement alone (*a* =1), we observe that the synaptic gating ratio, 1 − *c*, quickly approaches 0.5 (i.e., 50%) as the number of contexts increases (Fig. 7(a)). This trend also holds for synaptic refinement in combination with random neuronal gating (Fig. 7(b)) for parameter ranges with high overlap between contexts (low neuronal gating, 1 − *a*, and large number of contexts, *s*). The intuition for this is straightforward - when enough noise has been added to the weight matrix, each element has the desired sign for approximately half of the connections. Notably, this also implies that the number of contexts (and thus patterns) that the network stores can potentially be increased indefinitely, without affecting the fact that approximately half of the synapses will still reflect the appropriate correlation structure of the accessible memory patterns. We compare our results with simulations of weight matrices of *N* = 10,000 and confirm that our approximation for the overall connectivity holds (Fig. 7(c)).

**Fig. 7:**
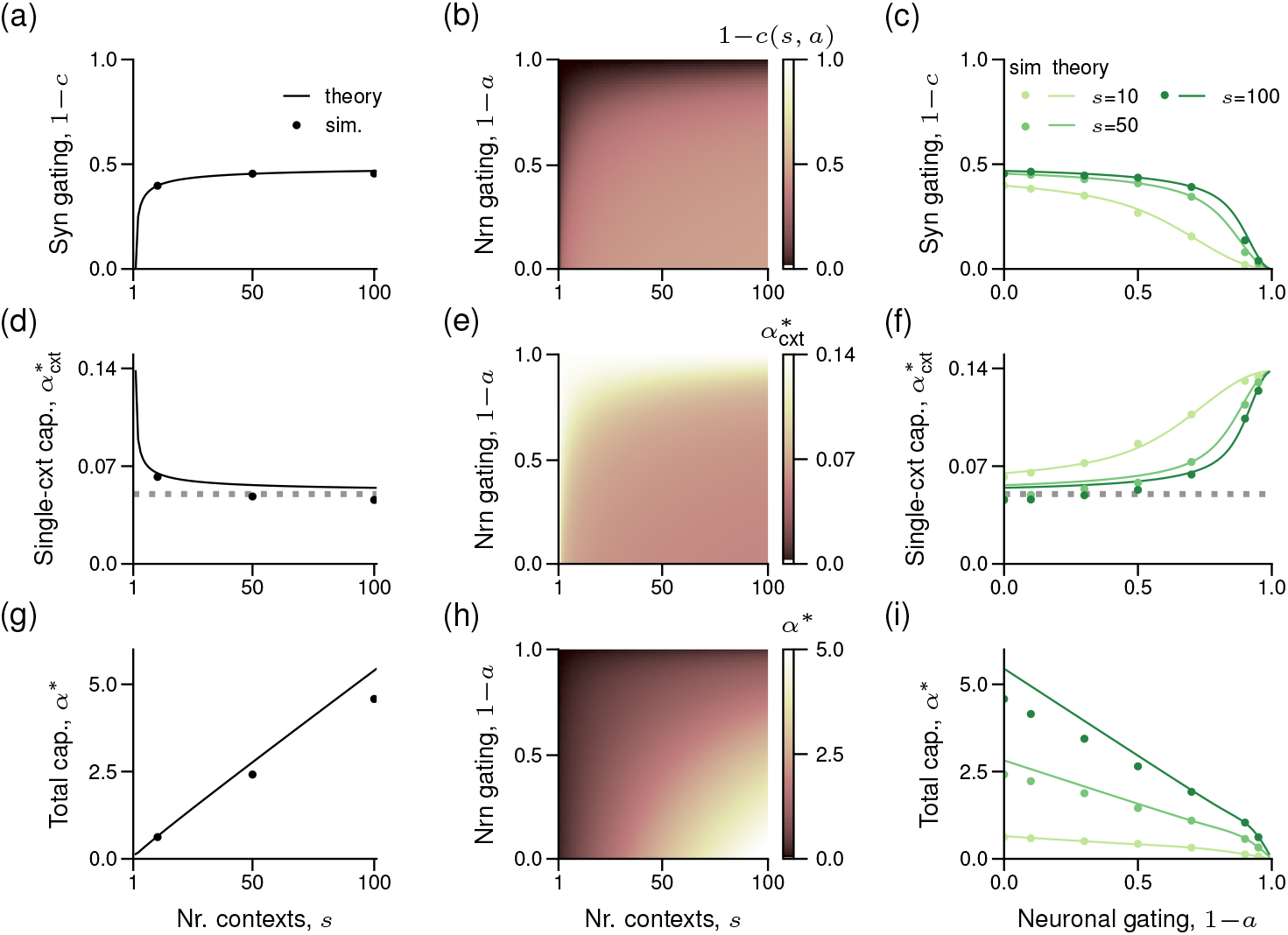
Synaptic gating ratio and memory capacity following synaptic refinement. (a-c) Estimation of the resulting synaptic gating ratio, 1 − *c* (Eq. 33), following synaptic refinement alone (panel (a)) or synaptic refinement with random neuronal gating (panels (b,c)). Synaptic gating ratio as a function of: number of contexts, *s* (panel (a)); number of contexts, *s*, and neuronal gating, 1 − *a* (panel (b)); and neuronal gating, 1 − *a* (panel (c)). (d-f) Estimation of single-context capacity, 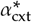 (Eq. 35), for synaptic refinement alone (panel (d)) or synaptic refinement with random neuronal gating (panels (e,f)). Single-context capacity as a function of: number of contexts, *s* (panel (d)); number of contexts, *s*, and neuronal gating, 1 − *a* (panel (e)); and neuronal gating, 1 − *a* (panel (f)). (g-i) Total capacity estimation, 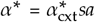 (Eq. 35), for synaptic refinement alone (panel (g)) or synaptic refinement with random neuronal gating (panels (h,i)). Total capacity as a function of: number of contexts, *s* (panel (g)); number of contexts, *s*, and neuronal gating, 1 − *a*, (panel (h)); and neuronal gating, 1 − *a* (panel (i)). Comparison with numerical simulations (dots) for three different numbers of contexts in panels (a,c,d,f,g,i): *s* = 10, *s* = 50, and *s* = 100. All numerical simulations from panels (a,c,d,f,g,i) were done with *N* = 10,000 (Appendix G).

### 3.2. Heuristic estimation of storage capacity and information content

To estimate the memory capacity with synaptic refinement, we need to take into account optimized synaptic gating. Synaptic refinement introduces correlations between the two crosstalk terms in Eq. 10, i.e., the within-context crosstalk and across-context crosstalk are no longer independent. We observe that, when neuronal gating is high (1 − *a* > 0.9), few connections are gated, and the single-context capacity is close to that of the standard Hopfield network 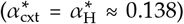. For lower amounts of neuronal gating (e.g., 1 − *a* < 0.5), synaptic gating rises to 0.5. In this regime, a given synapse is either zero (with ~ 50% probability) or assumes a value with the same sign as the correlation between all patterns in the active context, i.e., the sign of that synapse for a weight matrix of the active context (Eq. 31). However, the value of a given synapse may not reflect the actual correlation between the patterns in the active context, due to the noise added by the patterns stored in all inactive contexts. Such a network may thus be well approximated by a Hopfield network with binary weights^2,32^, but also scaled roughly linearly by sparsity^27^. The connections of a Hopfield network with binary weights follows a standard Hebbian learning rule that is passed through a sign-function, and weights are set to +1 or −1. In this case, the single-context capacity of a context-modular memory network can be approximated as

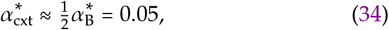

where 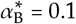 isthememory capacity of a network with binary synapses^32^. Therefore, to a rough approximation, we may be able to estimate the capacity in between two extremes – no synaptic gating, *c*(*s*, *a*) ≈ 1, and half of connections gated-off, *c*(*s*,*a*) ≈ 0.5 – by linearly interpolating between *α*_H_ and 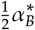. This interpolated estimation can be written as

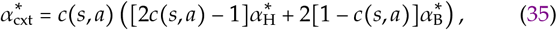

where 1 − *c*(*s*, *a*) is the synaptic gating ration defined by Eq. 33.

Based on this approximation, we observe that single-context capacity for synaptic refinement alone (*a* =1) is upper-bounded by the Hopfield limit, 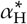, decreasing as the number of contexts increases (Fig. 7(d)), and following similar trends as for random neuronal gating alone for different neuronal gating ratios, *a* < 1 (Fig. 7(e,f)). However, this decrease in single-context capacity appears to have a lower bound as well, near 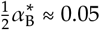 (Fig. 7(d,f), dashed gray line), as predicted by the estimated theoretical capacity (Eq. 35). This finite lower bound leads to a total capacity that increases roughly linearly with the number of contexts, *s*, and decreases roughly linearly with neuronal gating ratio, 1 − *a* (Fig. 7(g,h,i)), growing to large values (*α*\* ≈ 5 for *s* = 100 contexts) – up to 30 times that of the standard Hopfield network (Fig. 7(g)). Numerical simulations confirmed the theory both for synaptic refinement alone (Fig. 7(a,d,g)) and in combination with random neuronal gating (Fig. 7(c,f,i)). For both the theoretical and numerical results, we observe that the optimal neuronal gating is zero, i.e., *a*_opt_ = 1, and so the addition of neuronal gating to synaptic refinement does not further increase memory capacity.

The information content of the context-modular memory network with synaptic refinement grows to much larger values than both standard and low-activity Hopfield networks. Writing the information content as a function of the information content of a standard Hopfield network, we arrive at 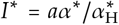. This means that the information content grows with the total capacity, *α*\*. For example, for *a* = 1 and *s* = 100,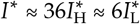. For the total capacity, *α*\*, and the total information context, *I*\*, to be high, the contextual control requires that half of synapses to be gated off for each context. However, this type of control requires many more additional neurons for autonomous implementation, which decreases the overall information content (see Discussion).

### 3.3. Arbitrary allocation of memories with synaptic refinement

Synaptic refinement is only applied to the network’s recall dynamics, and thus memories do not have a strict grouping as they did for random gating. We therefore momentarily take a different perspective on context-dependent memory, in which a base set of memories is stored in the weight matrix, and then arbitrary memory subsets may be stabilized in different contexts at the time of recall. To illustrate this feature, we first store *P* memory patterns without explicitly specifying contextual states,

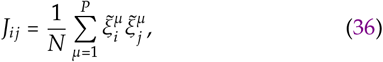

where 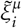 (*i* = 1…*N* and *μ* = 1… *P*) are context-independent memory patterns defined by 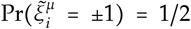. Next, we assign an arbitrary group of these memories, of size *p*, to be part of a context *σ*, and determine the appropriate contextual configurations, i.e., the set of gating variables 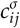 that enables successful memory recall (Eq. 32). Importantly, this is only possible because all neurons remain ungated in each context, even if individual connections are gated off, which means that *a* = 1.

We observe that arbitrary sets of memory patterns can be reliably retrieved (Fig. 8(a)), provided that the number of patterns stored in the active context is sufficiently small (*α*_cxt_ ≤ 0.03N). Performance begins to degrade when the number of patterns per context approaches the single-context capacity, 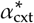, which depends on the overall load, *α*. Intuitively, the single-context capacity, 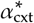, decreases with the increase in total load, *α* (Fig. 8(b)). As before, synaptic refinement increases the number of gated synapses during recall of memories in the active context as the total memory load is increased (not shown). This indicates that synaptic refinement can be used at any moment in time after memories are learned to stabilize an arbitrary group of memories, even if some of these memories are also allocated to other contexts.

**Fig. 8:**
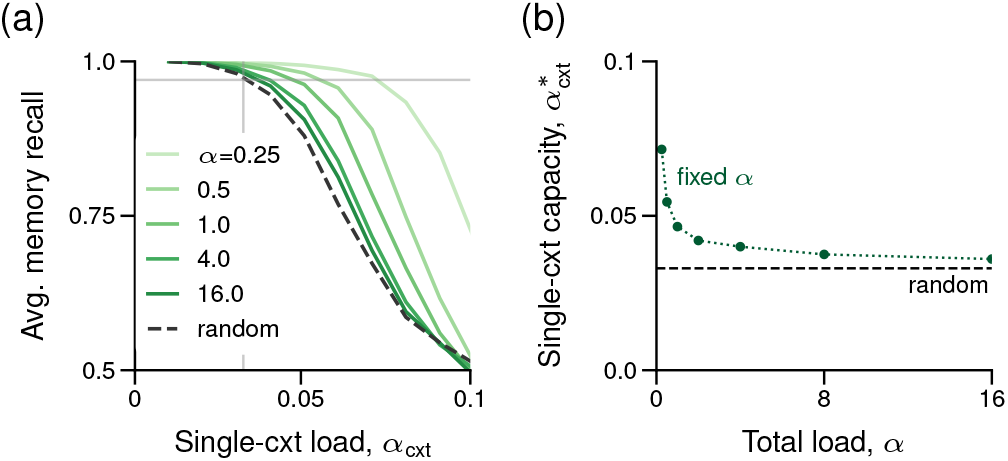
Arbitrary memory allocation through synaptic refinement. (a) Network performance as measured numerically by the average overlap between initial pattern and steady-state across all patterns for the active context, plotted for various total loads, a, as a function of the single-context load, *α*_cxt_. Simulation with a random weight matrix is shown by the grey dashed line. *N* = 1000 for all simulations. (b) Single-context capacity, 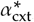, as a function of total load, *α*, estimated numerically from the simulations shown in panel (a). Single-context capacity for the random weight matrix is plotted as the dashed black line, also estimated from panel (a).

### 3.4. Stabilization of memory patterns in a random weight matrix with synaptic refinement

Given that synaptic gating modifies the energy landscape through a set of gating variables, even a randomly defined weight matrix may be utilized to recall memory patterns as long as the correct synaptic refinement is imposed. As a proof-of-concept, we thus generate a random weight matrix given by

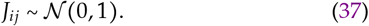

We then construct a set of *p* memory patterns, 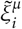 (*i* = 1…*N* and *μ* = 1… *p*), and define the gating variables 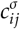 for the active context *σ* following Eq. 32. Remarkably, a random weight matrix can be successfully used to represent stable context-dependent memories (Fig. 8, dashed lines), as long as the single-context load is small (*α*_cxt_ ≤ 0.03).

Intuitively, this is possible because a random weight matrix will have, on average, half of its elements with the correct sign according to the correlation structure of the desired subset of patterns. This means that synaptic refinement can be used to impose any activity pattern as a stable attractor independent of the weight matrix, i.e., groups of memories are stored in the synaptic gating structure. Note that this requires the weight matrix to have a minimal amount of structure (e.g., randomly distributed with zero mean). This suggests that synaptic weights can be corrupted substantially without affecting performance, as long as the synaptic gating structure remains intact. Furthermore, it enables reliable recall with bounded (e.g., binary) synapses, an important consideration for biological memory^33^. In other words, by precisely controlling the synaptic gating variables, it is possible to recall a set of memories from a weight matrix that doesn’t contain any information about those memories.

## 4. Accessibility (stability)

A crucial feature of the model is the assumption that only a subset of memories need to be stable in each context. While storage capacity provides an indirect measure of memory stability for accessible memory patterns, our model also assumes that inaccessible memories are hidden, and thus unstable, in contexts for which they do not belong. Furthermore, as the number of contexts, *s*, increases, the proportion of accessible memories decreases as 1/*s* (Fig. 9(a)), leading to a clear trade-off between the storage capacity gains (which increases with s), and memory accessibility across all patterns at any given time. This also means that the proportion of inaccessible memories grows with the number of contexts, making it just as important to ensure that the hidden memories are unstable, as it is to ensure that the accessible memories are stable. To confirm that accessible memories remain stable while inaccessible memories remain unstable, we measured the stability of all memories.

**Fig. 9:**
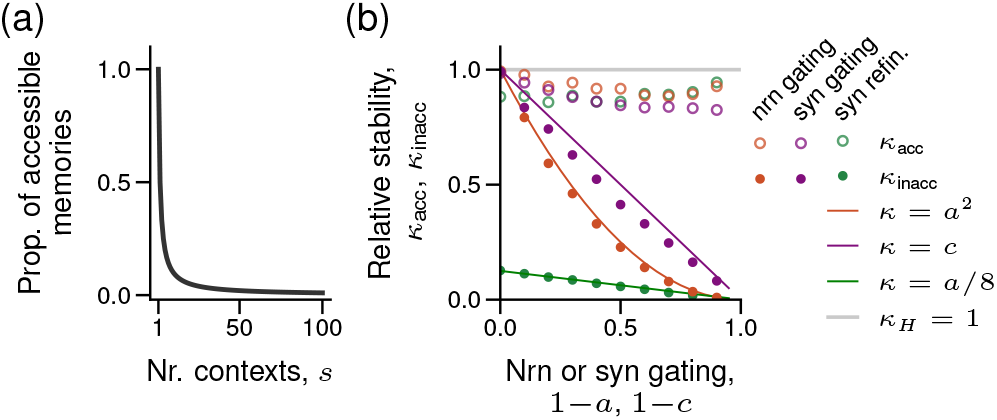
Memory accessibility. (a) Proportion of accessible memories, i.e., ratio of memories stored per context divided by the total number of memories, 1/*s*, as a function of the number of contexts, *s*. (b) Average memory stability for accessible memories, 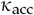 (Eq. 40, open circles), and inaccessible memories, 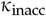 (Eq. 41, closed circles), with neuronal gating alone (red circles), synaptic gating alone (purple circles), or neuronal gating with synaptic refinement (green circles). Three comparison functions (full lines) are described in the legend. Stabilities are plotted relative to that of the standard Hopfield network 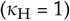.

To do so, we measure the so-called *stability parameter* of each memory pattern^34^, which provides a rough measure of the size of each pattern’s basin of attraction. We write the stability of a particular pattern *v* of context *σ* (for an arbitrary neuron i) when a (potentially) different context *k* is active as

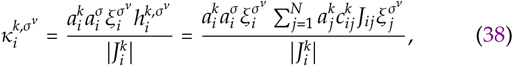

where 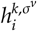 denotes the input to unit *i* when context *k* is active but the network is representing pattern *v* of context *σ*, and

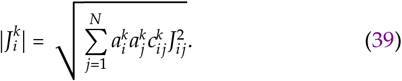

Averaging over all neurons, patterns, and contexts, we obtain an average stability parameter of accessible memories when setting *k* = *σ* in Eq. 38,

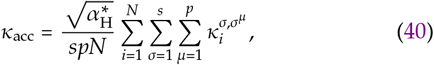

where we include the factor 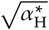 to make the measure relative to the stability of the standard Hopfield network (with mean stability 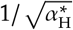)^34^. We can then also measure the stability of inaccessible memory patterns by setting *k* ≠ *σ* in Eq. 38. As above, averaging over all neurons, patterns, and contexts we obtain an average stability parameter of inaccessible memories,

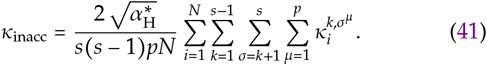

We measured the stability of both accessible and inaccessible memories at full capacity (*N* = 10,000, see Appendix G). For random neuronal and synaptic gating, as well as synaptic refinement, we observe that the stability of accessible memories is close to the stability levels of the standard Hopfield network and independent of the number of contexts, *s*, as well as neuronal and synaptic gating ratios, 1 − *a* and 1 − *c* (Fig. 9(b), open circles). As for inaccessible memory patterns, we find that their average stability is always equal to or lower than that of accessible memory patterns (Fig. 9(b), filled circles). The average stability of inaccessible memory patterns depends upon gating ratios, but remains relatively independent of the number of contexts, *s* (not shown). For neuronal gating, the stability of inaccessible memories increases roughly with *a*^2^ (Fig. 9(b), orange line), whereas for synaptic gating, the stability of inaccessible memories increases with *c* (Fig. 9(b), purple line). For synaptic refinement with random neuronal gating, the stability of inaccessible memories decreases as the number of contexts, *s*, increases (not shown), because of the dependency of the synaptic gating ratio on the number of contexts. In the limit as the gating ratio increases to 50% (i.e., for *s* > 100), the stability of inaccessible memories remains very low for all values of *a*, and is well approximated by *a*/8 (Fig. 9(b), green line). Overall, we conclude that the modular organization of memories successfully suppresses inaccessible memories while maintaining high stability of accessible memories, as intended.

## Discussion

The growing consensus perspective in neuroscience and psychology views biological memory not as a static storage device, but rather as a dynamic and flexible system^20,24^. Such a system adapts to current behavioral states, i.e., contexts, and is thus able to modify the understanding of past experiences when needed^35^. Here, we revisited classic theoretical work on auto-associative memory in light of this perspective. Specifically, we studied a model in which the memory patterns and network connectivity are modified through neuronal and synaptic gating by a background contextual state. We found that such a scheme not only results in increased memory capacity, but is also effective at modulating the stability of stored patterns, thereby selecting a subset of memories for recall and reducing interference from others. Improved memory capacity is thus not the main feature of the context-modular memory network, but a side-effect of increased control and flexibility which is traded off with the proportion of accessible memories at any given time.

We explored two contextual gating schemes: neuronal and synaptic. To implement these schemes, an inhibitory mask is imposed onto the recurrent connectivity to modulate the energy function of the network dynamics. Neuronal gating acts to remove particular neurons from the active context, thereby shrinking the effective network size and removing interference from masked neurons. The potential biological mechanisms for neuronal gating are numerous, including peri-somatic inhibition^36^ and modulation of intrinsic excitability^37^. Synaptic gating features the masking of individual synaptic elements, thereby making the network connectivity more sparse (diluted) in a particular context, reducing interference between accessible and inaccessible memory patterns. Given the precise and complex nature of synaptic gating, it is likely that full individual synaptic control is not plausible (but see ref.38). However, there is considerable evidence for control of synaptic inputs at the level of dendritic branches^39^, supported by findings such as dendritic clustering^40^, localized dendritic inhibition^41^, and localized neuromodulation^42^. One area for future work would be to relax the precision of synaptic gating, and explore a continuum between full individual control and coarse control of all inputs (as in neuronal gating).

We proposed two methods for setting contextual states in our model: random allocation and refinement. In random gating, contextual states are imposed at the time of learning, which pre-allocates neurons and synapses to store memory patterns in a given context, and does not affect gated components. For random gating to be effective, contextual states are also re-instated during recall — such contextual reinstatement is hypothesized to occur in the brain^43,44^. We observed that neuronal gating was much more effective than synaptic gating for random allocation, resulting in high memory capacity (Fig. 2) and low stability of inaccessible memories (Fig. 9). Such a finding is intriguing in light of recent evidence for allocation of particular neurons (through excitability changes) to memories in the amygdala and hippocampal system^26,37^. Specifically, the activity levels observed in such studies (~ 20 − 30%) correspond to optimal neuronal gating levels in our model for tens to hundreds of contexts (Fig. 3).

The second contextual gating method proposed was refinement, in which gating states are optimized to strengthen accessible memories after learning (i.e., contextual states are imposed only at retrieval). This scenario was proposed specifically with synaptic control in mind, as synapses were gated off in a particular context if they had a net harmful effect on recall (refinement of neuronal activity would result in altered memory patterns, so it was not considered). We found that synaptic refinement resulted in even larger gains in memory capacity (Fig. 7) and better control of memory accessibility (Fig. 9), with connectivity levels approaching 50% (Fig. 7). Such a scheme fits well with the idea that memories are constantly being updated, with each recall event possibly leading to changes^35^. Furthermore, recent work has also suggested that memory expression can be masked and controlled through inhibition^45,46^. Finally, the finding that synaptic refinement can stabilize arbitrary groups of memory patterns (even using a random weight matrix) suggests that the precision of synaptic learning may not be crucial to long-term maintenance of memories (see refs. 33,47,48 for related studies).

Memory capacity, stability, and information content results were obtained using conditional measures — i.e., we assumed that context is externally controlled, and memory stability is measured conditioned upon this fixed state. This choice was made due to reasons of analytical tractability, and the desire to model a flexible memory system more related to biological memory^20,24^. Furthermore, we contend that the questions addressed here do not depend upon an explicitly-specified context representation, as the focus was rather on the consequences of contextual control for an associative memory network. However, our conditional measures make comparisons with previous capacity estimates difficult, as performance metrics usually require stabilizing all memories at all times^2^. We therefore stress that our aim is not to compare the capacity with previous networks, but rather to propose a framework within which to study capacity while emphasizing flexibility and accessibility. In future work, an explicit implementation of a context-encoding network may potentially lead to interesting biological comparisons, including dynamic competition between memories^25^, as well as descriptions of pathological memory states like in amnesia^49^. Overall, there may be a tradeoff between capacity and flexibility, and it may be in the interest of the brain to favor flexibility over capacity. Theoretically, it will be necessary to develop richer measures than capacity to determine the use and function of memory that includes flexibility, retrieval dynamics, and evolving memory patterns^50–52^.

One simple way of considering the complexity of such contextual representations in the capacity estimations is to consider a Hopfield-like (context-encoding) network storing contextual states as memory patterns. For neuronal gating, our model would require the size of such a network to scale with the number of contexts, *s*, which should be relatively small compared to the memory network size, and thus result in qualitatively similar results. For synaptic gating, however, our model would require fundamentally different dynamics in which synaptic weights are controlled by activities of other neurons^53^. Assuming this is possible, and constraining each context-encoding neuron to have on the order of *N* synapses, the network should scale with *sN*, leading to a context network that is potentially orders of magnitude larger than memory network, and reducing the memory capacity below the Hopfield limit with only a few contexts. Such intuitions argue against full and precise synaptic control. However, relaxations of such precise control, especially for a smaller number of contexts, may be possible, and our model serves as a proof of concept of such a situation.

Context-modular memory networks utilize several principles which are common in previous associative memory models, including sparsity, modularity, and hierarchical organization^2,10^. Modularity can be used to mimic the architecture of the cortex –; dense local with sparse long-ranged connectivity^12–14^ – and several other models have explored a hierarchical organization of memories^8^, or non-tree-like organization of semantic memories^54^. Our model bears a resemblance to these previous models, but is unique in considering context as the main determinant of memory groupings (but see ref.55), as well as modeling context as an external signal imposed on the memory network (in contrast to previous hierarchical models). Extensions to associative memory networks that improve memory capacity, such as low activity patterns^6^, alternative learning rules^19^, or complex synapses^56^ may be incorporated into the context-modular memory network architecture for additional benefits. Finally, other models have considered contextual control of learning and memory^57–60^, as well as similar gating mechanisms applied in other situations (e.g., cortical information routing^61–63^, deep learning^64^, inhibitory learning^65^, and motor learning^66^), highlighting the importance of contextual flexibility for cognition.

In conclusion, the integration of associative memory models with retrieval processes and decision making is a promising area of future research^67^. Organizing memories around context leads to control over memory access, which the brain may harness. We propose the contextual gating framework introduced here as a promising model for flexible memory access which may lead to new insights into biological and artificial memory.

## Acknowledgements

We thank Helen Barron, Vezha Boboeva, Adam Packer, João Sacramento, Andrew Saxe, Misha Tsodyks, and Friedemann Zenke for helpful comments at various stages of this work, and Rubem Erichsen Jr for carefully reading the manuscript and valuable comments. This work was supported by a Sir Henry Dale Fellowship by the Wellcome Trust and the Royal Society (WT100000; WFP, EJA and TPV), a Wellcome Trust Senior Research Fellowship (214316/Z/18/Z; EJA and TPV), and a Research Project Grant by the Leverhulme Trust (RPG-2016-446; EJA). For the purpose of open access, the authors have applied a CC BY public copyright license to any authors accepted manuscript version arising from this submission.

### Appendix A: Equivalence of {±1} and {0,1} formulations

We define the context-modular memory network considering three states, 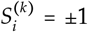 for neuron *i* allocated to the active context *k* and 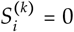 for neuron *i* gated off in the active context *k*. This formulation can be modified to a {0,1} formulation, making it effectively a network of binary neurons. To see this, it is sufficient to show that the dynamics in either case are equivalent. In the {0,1} formulation^2^, we consider the state of neuron *i* as 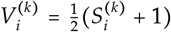 for neuron *i* allocated to the active context *k* and 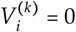 for neuron *i* gated off in the inactive context, and memory patterns given by 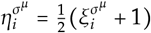. From these definitions, and considering a neuron *i* in the active context *k*, we arrive at

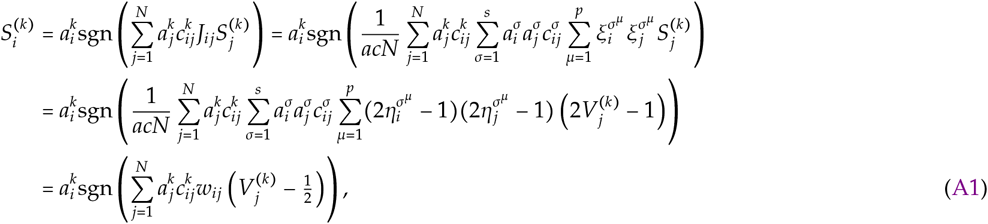

and thus the input to neuron *i* can equivalently be given in terms of *V* variables. We can thus write

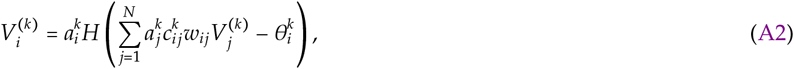

where *H*(·) is the Heaviside function, such that 2*H*(*x*) − 1 = sgn(*x*), *w_ij_* is the weight matrix, and 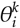 is a context-dependent threshold. In this formulation, the weight matrix, *w_ij_*, is given by

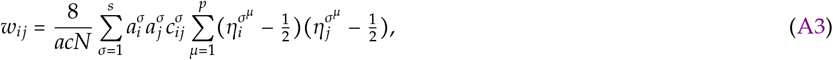

and the (context-specific) threshold 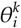 is

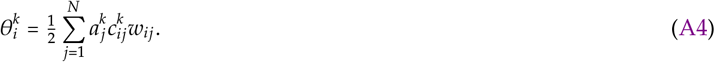

Thus, the deterministic dynamics of the two formulations are equivalent. We note that this holds also for the stochastic version of the dynamics, in which *V_i_* = *σ*(*h_i_*) = *e*^2*h*_i_^/(*e*^2*h*_i_^ + 1) and *S_i_* = tan(*h_i_*), due to the identity tanh(*x*) = 2*σ*(*x*) − 1.

### Appendix B: Wald’s equation

For the signal-to-noise analysis (Section 2.1), we make use of a result in statistics known as Wald’s equation^68^, which we summarize here. Consider a binomial random variable 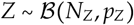. Let 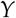 be the sum of a sequence of independent identically distributed random variables *X*_i_ of length *Z*, i.e., 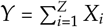. Assuming that each *X*_i_ is independent of *Z*, then the mean and variance of 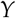 can then be written as

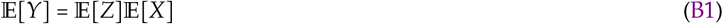

and

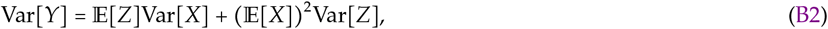

where we have dropped the index on *X* because each random variable *X*_i_ comes from an identical distribution.

### Appendix C: The standard, diluted, and low-activity Hopfield models

We compare the results obtained in this study with that of the standard Hopfield model^1^. For a network of *N* neurons storing *p* patterns, we use *α*_H_ = *p*/*N* to denote the load (with 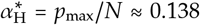 at capacity) and *m* to denote the overlap of the state configuration with the retrieved pattern (*m*\* ≈ 0.97 at capacity)^18,30^. For convenience of the reader, we write the three mean field equations for the standard Hopfield network here, as

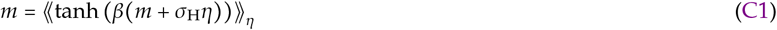

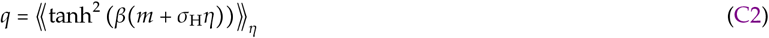

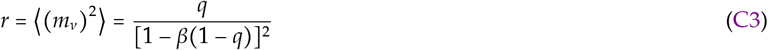

where 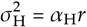 is the crosstalk noise and 《·》_*η*_ represents a Gaussian average, i.e., 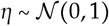.

Next, we utilize previous results for a diluted Hopfield network^27^ with dilution parameter *c* (analogous to the synaptic gating parameter in this work). The diluted connectivity was shown to be well-approximated by Gaussian noise^27^,

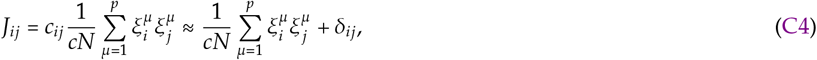

with

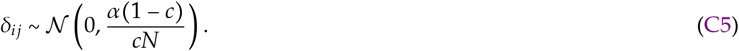

This weight matrix results in analogous mean field equations as the Hopfield network (Eqs. C1–C3), but with a noise term

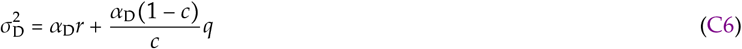

in place of *σ*_H_. This results in a capacity that scales roughly linearly with 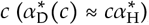, and an overlap that also degrades as *c* becomes small (Fig. 4, purple line). Using the mean-field results, it can also be shown that a signal-to-noise analysis of a diluted Hopfield network results in an input (to neuron *i* for a pattern *v*) of

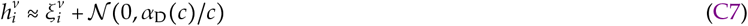

which we utilize in Section 2.1.

Finally, we consider an associative memory variant with low-activity patterns^6^ (Fig. 3(b) and Section 2.3). The capacity of the low-activity Hopfield network, 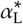, scales with the activity level as

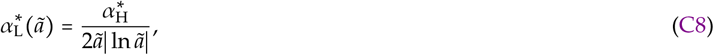

for *ã* ≪ 1^6^ (the 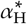 factor is included to achieve a more fair comparison for more intermediate *ã* values).

### Appendix D: Mean-field derivations

We used a heuristic mean-field approach^2,69,70^ to compute the capacity in Section 2.2, which we detail here. Following ref.27 (Eq. C4), we simplify the weight matrix used in the random neuronal and synaptic gating scheme (Eq. 5) using an effective matrix, 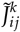 so that dilution is not explicitly included,

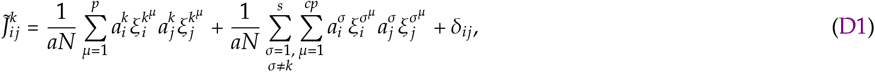

where *δ_ij_* represents the effects of dilution within context *k*,

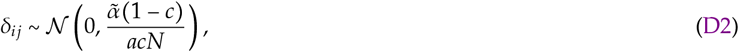

and 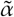 is the effective noise in the weights due to all stored patterns, which we leave unspecified for now. Synaptic gating reduces the original sum over (*s* − 1)p terms to an average of *c*(*s* − 1)*p* – the second term of the right-hand side of Eq. D1 − which we take into account by changing the limit of the sum over patterns to *cp*.

We describe the dynamics of the system using three pairs of order parameters. First, we use the two load parameters as defined in Eqs. 3 and 4 (single-context and total load). Second, considering that the currently active context is *k*, we use 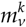 and 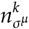 to denote the *overlap* between the average network state 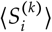 and a particular pattern *v* of context *k*, 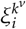, and a pattern *μ* of context *σ*, 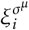, respectively,

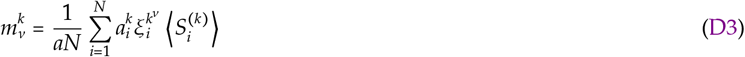

and

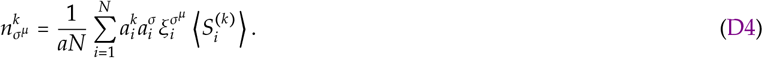

Third, we use *r^k^* and 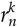 to denote the *mean square overlap* of the system configuration with the nonretrieved patterns in context *k*, and all other contexts, respectively,

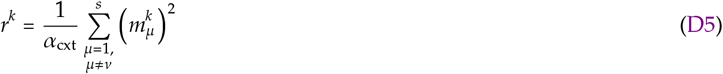

and

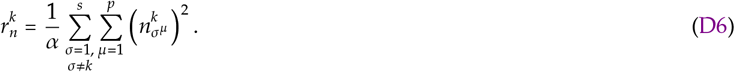

We consider that the network state is close to pattern 1 of context *k*. This means that if this pattern is a fixed-point of the network’s dynamics when context *k* is active, 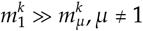.

We then follow analogous steps outlined in refs. 2,69,70 by plugging the mean field equations for 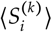 from Eq. 20 into Eq. D3, and expanding the weight matrix using the expression in Eq. D1,

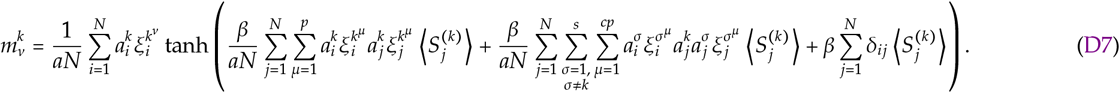

We then rearrange terms and substitute the other 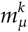 and 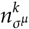 terms into the equation to obtain

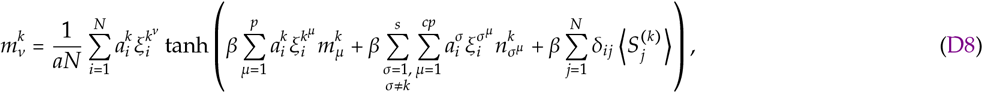

and then separating terms and multiplying by 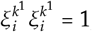, we get

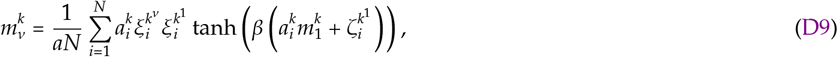

where we use 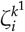 to denote the crosstalk terms,

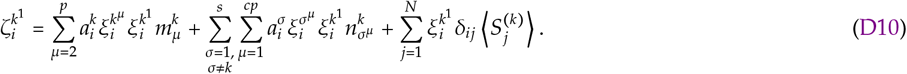

We then repeat these steps twice more (plugging Eqs. D1 and 20 into Eq. D3) — once using 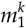 (the retrieved pattern) in place of an arbitrary 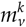, and another for 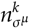 (using Eq. D4 in place of Eq. D3), to obtain

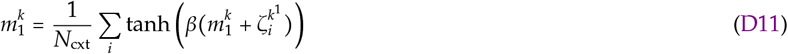

and

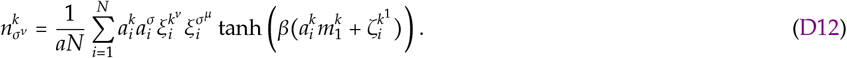

These expressions are analogous to that of the standard Hopfield network and we refer the reader to refs. 2,69,70 for the details involved in manipulating these expressions to obtain the final mean field equations. One final detail unique to the context-modular memory network is estimating the variance of the crosstalk noise in Eq. D10. Due to the similarity with the diluted Hopfield network (Appendix C), we can write the standard deviation of the term as

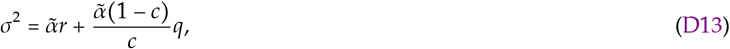

with 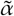 being the effective load of the network due to the first two terms in Eq. D10. Note that the second term comes from the definition of *δ_ij_* above (Eq. D2). Following similar arguments to the signal to noise analysis, we expect the effective load to depend on a particular realization of the number of overlapping contexts,

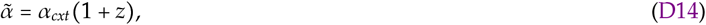

where *z* is a binomial random variable distributed as 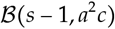 (see Eq. 13) and with expected value (*s* − 1)*a*^2^*c* (see Eq. 17).

Finally, from Eqs. D9, D11 and D12, we obtain the three mean-field equations

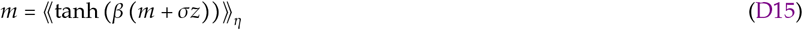

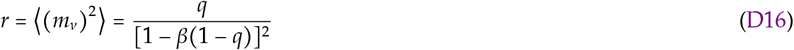

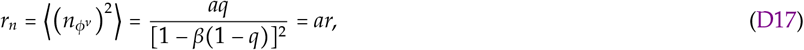

where *q* is defined as

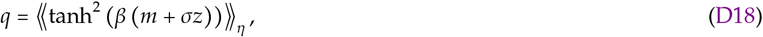

and 《·》_*η*_ represents a Gaussian average 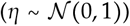, and σis defined in Eq. D13. Similar to the signal to noise analysis, the binomial random variable *z* can be dealt with by performing a binomial average at the end, or by approximating it with the expected value.

These equations can be solved numerically using standard methods. We again take a shortcut here and note the correspondence to the standard and diluted Hopfield networks, enabling us to avoid numerically solving the mean field equations. Following Section 2.1, we can set the variance term here equal to that of previous results, and obtain Eqs. 15 and 16. By simplifying the binomial random variable *z* to the expected value, we obtain Eqs. 17 and 18. Finally, we can write the mean field overlap with pattern 1 of context *k* as *m*(*c*, *a*, *s*) = *m*_D_(*c*) as written in the main text.

### Appendix E: Information content

Information content can be estimated based on the entropy (Shannon information) across all retrieved patterns^19^. Importantly, we consider that the total magnetization is always zero in our calculations — i.e., for both standard and diluted Hopfield networks, we consider that half of the neurons assume value +1, while half assume value −1. We can write the entropy per neuron considering the original patterns as

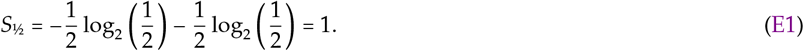

The entropy per neuron considering the missing information and average overlap, *m*, is given by

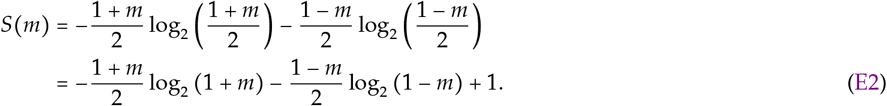

Combining Eqs. E1 and E2 we arrive at the total information content for the diluted Hopfield network as

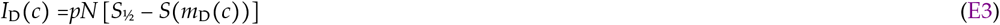

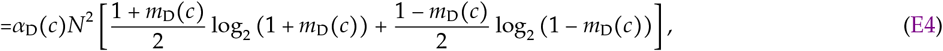

with *I*_H_ = *I*_D_(*c* = 1) being the information content of the standard Hopfield network.

Using the same argument, the maximum information content of the low-activity Hopfield network, when the activity level goes to zero, can be written as

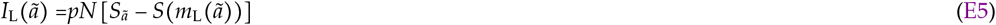

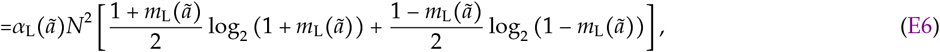

where *S_ã_* = *ã*log_2_*ã* − (1 − *ã*) log_2_(1 − *ã*) is the entropy per neuron for an activity level *ã* and *m*_L_(*ã*) is the average overlap between original patterns and minima of the network’s dynamics for activity level *a.* For the limit of vanishing activity level, we get^19^

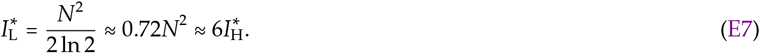

For the context-modular memory network, we consider the conditional information content of patterns stored in networks of size *N*_cxt_. This way we can rewrite Eq. E4 substituting *α*_D_(*c*)*N*^2^ by *αaN*^2^, arriving in Eq. 29.

### Appendix F: Connectivity density for synaptic refinement

The expected synaptic gating ratio, 1 − *c*(*s,a*), for synaptic refinement with random neuronal gating (with parameter *a*), can be estimated assuming that the weight distributions for a single context and across all contexts are Gaussian with zero mean and variances proportional to the amount of crosstalk that they contribute. Given this, the probability that a particular weight will be removed can be approximated as

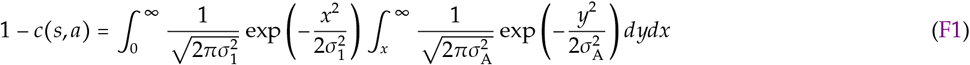

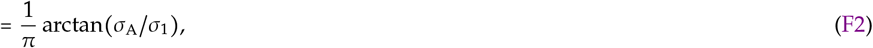

where *σ*1 and *σ*_A_ are the crosstalk standard deviations from patterns within the same context and all other contexts, respectively, whose ratio is

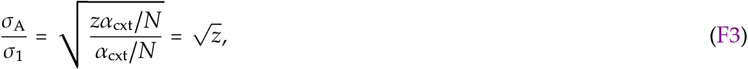

and *z* being the binomial random variable distributed as 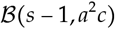. Doing a binomial average, we obtain the same equation in the main text,

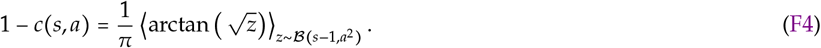

### Appendix G: Details of numerical simulations

We briefly describe the details of all numerical simulations here, used to generate the overlap and capacity plots in Figs. 4,5,7, and 8. Code was written in python with the help of Pytorch^71^. Memory capacity was estimated numerically by building finite-sized networks (N =10,000) initialized with a set of random patterns and connectivity as defined in the main text. To test stability of the patterns, we initialized the network in each memory state and simulated the dynamics until they either reached a steady-state or they reached the maximum number of allowed time steps (60). Dynamics were run *synchronously* (i.e., all units were updated simultaneously) according to Eq. 1, and recall performance was measured using the overlap

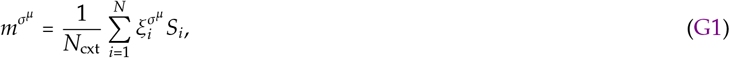

where 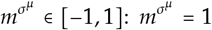 when the state is exactly the pattern 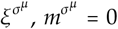 when it is uncorrelated with the pattern 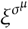, and 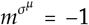 when the state is the exact opposite (anti-correlated) of pattern 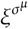. We repeated this procedure for all patterns in a particular context to obtain an average overlap 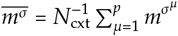. Capacity was set as the maximum number of patterns stored for which the average overlap was greater than or equal to the overlap curves in Fig. 4 (as a function of gating ratios 1 − *a* and 1 − *c*). Synchronous dynamics were chosen for efficiency reasons despite potential convergence issues. To generate the overlap plots in Fig. 4, we simulated networks with the theoretically-expected number of patterns at capacity (using dynamics as specified above), and measured the resulting average overlap. All code used to run simulations and generate the paper figures will be made available upon publication of this work.

